# High-throughput and in-depth analysis of plasma proteome by UV-assisted protein digestion, 5-plex labeling and MS-based high abundance protein removal method

**DOI:** 10.1101/2023.10.20.563193

**Authors:** Zhiting Wang, Boxin Qu, Weijie Zhang, Zhen Liang, Liming Wang, Jianhui Liu, Huiming Yuan, Lihua Zhang, Yukui Zhang

## Abstract

Human plasma contains various proteins, some of which reflect individuals’ physiological health state or predict diseases. Therefore, plasma analysis can provide a wealth of information on novel biomarkers for clinical diagnosis and prognosis. The use of mass spectrometry (MS) for high-throughput and in-depth quantitative proteomic analysis of plasma allows for highly specific and quantitative readout, but is challenging because of the high dynamic range of protein abundances. Here, we introduce a robust, high-throughput, and in-depth workflow for plasma proteome analysis based on UV-assisted rapid protein digestion, 5-plex dimethyl labeling, and MS-based high abundance protein removal. UV-assisted protein digestion could quantify the same protein numbers as traditional enzymatic hydrolysis and achieve a low miss-cleavage rate in only 20 minutes. And the MS-based high abundance protein removed 5-plex DIA method, which does not require extra protein depletion procedures, enables quantitative analysis of more than 600 proteins using an equivalent MS analysis time of 30 minutes per sample. The average relative error was 6.9%. We believe the method is beneficial for analyzing large numbers of trace-level clinical samples and broaden a vision for the discovery of low abundance protein markers.

## 1. Introduction

Plasma is a gold mine which contains a large number of secreted or leaked proteins from tissues, the concentrations are highly influenced by individual physiological and pathological conditions. Therefore, the analysis of plasma proteome can provide substantial information on novel biomarkers for diagnosis, prognosis and personalized medicine. However, plasma is a complex biochemical matrix whose proteome spans a dynamic range in excess of 10 orders of magnitude in which the 10 most abundant proteins account for 90% of its total protein mass^[1]^. The considerable complexity and large dynamic range of protein concentrations present substantial analytical challenges, especially for large cohorts. Therefore, although plasma is an ideal window for reflecting health/disease status, only few novel biomarkers have been established in clinical practice^[2-4]^, improving the efficiency of the biomarker discovery process remains a critical need.

Depletion or enrichment strategies have been widely used for discovery workflows as they address the inherent mass imbalance and broad dynamic range of plasma^[5]^. Common depletion strategies such as immunoaffinity-based assay^[6]^ usually target at albumin, immunoglobulins, transferrin, fibrinogen and other high abundance proteins (HAPs). It has now become possible to identify more than 1,000^[7,8]^ or even more than 5,000 proteins^[9]^ in plasma. However, immune-depletion may result in the nonspecific depletion of protein subsets with which proteins targeted for depletion interact, adversely leading to quantitative biases^[10]^. To solve the protential low abundance proteins (LAPs) loss problem, magnetic nanoparticles (NPs)^[11,12]^ that can enrich LAPs have emerged. After the surface of NPs contacts plasma, a wide range of proteins can be adsorbed on the surface to form a “protein corona”^[13,14]^, achieving enrichment of low abundance proteins. This method has been used for the discovery of new plasma biomarkers due to its deep coverage, and more than 1000 proteins were quantified for ovarian, or lung cancer study^[12,15]^. However, the accuracy of plasma quantification after treatment with NPs still needs to be considered. Besides, these pre-treatments decrease throughput and increase expenses, which are undesirable in clinical practice.

Here, we sought to address these problems by developing a robust and highly streamlined plasma proteomics workflow. Firstly, we developed a rapid sample processing system based on UV-assisted protein digestion (UVAPD), which could achieve the depth of traditional enzymatic hydrolysis and a low miss-cleavage rate in 20 minutes. Secondly, we established a MS-based high abundance protein removed 5-plex DIA method. It does not need for extra sample procedures and could quantified over 600 proteins with dynamic range spanning 9 orders of magnitude using 0.5 h LC-MS analysis time per sample, with an average relative error of 6.9%. We hypothesized that the resulting workflow would have a high yield of information about individual health status and could be obtained for a large number of trace-level clinical samples.

## 2 Experimental section

### 2.1 Sample Preparation

Human blood was collected in blood collection tubes and spun-down 1500×g for 15 min in a clinical centrifuge to separate cells from the plasma then the plasma was aliquoted and stored at −80°C. The proteins extracted from plasma were denatured and reduced by 10 µM DTT at 95°C for 15 min, followed by protein digestion under 365 nm UV irradiation for 5 min with trypsin or a mixture of sequencing-grade trypsin and Lys-C (the mass ratio of 2:1). The mass ratio of protein to enzyme was set to be 25:1(m/m). For comparison, the same aliquot of proteins was also prepared by traditional in-solution digestion protocol. Briefly, the proteins from human plasma were denatured and reduced by 10 µM DTT in 95°C for 15 min, then alkylated by 20 µM IAA in the dark at room temperature for 40 min. Finally, the protein solution was digested by trypsin at 37°C overnight under the same mass ratio of protein to enzyme (30:1,m/m). The peptide concentration was determined by Nanodrop (Thermo, Rockford, USA).

### 2.2 Dimethyl labeling

The plasma digests were labeled by 28 (4% CH_2_O and 0.6 M NaBH_3_CN), 30 (4% CH_2_O and 0.6 M NaBD_3_CN), 32 (4% CD_2_O and 0.6 M NaBH_3_CN), 34 (4% CD_2_O and 0.6 M NaBD_3_CN) and 36 (4% ^13^CD_2_O and 0.6 M NaBD_3_CN) dimethyl labeling reagents, and incubated for 1h at room temperature. The 32 labeled digests were used for spectral establishment, otherwise the five labeled digests were mixed at the ratio of 1:1:1:1:1 for MS-based high abundance protein removed 5-plex DIA analysis. The above samples were desalted by homemade C18 column and stored at -20°C before usage.

### 2.3 High pH reverse-phase fractionation

High pH reverse-phase fractionation (HpHRPF) were performed to build an in-depth labeled plasma spectrum library by using a liquid chromatography (Agilent, Tokyo, Japan) with a homemade reverse phase column (C18, 5 μm, 100 A□, 2.1× 150 mm i.d., Durashell, Tianjin, China) under mobile phase at pH=10 (solvent A: 100% water, ammonia was added until pH reached 10.0; B: 80% ACN, same volume of ammonia was added to solvent B as A). Peptide samples were loaded to the column and separated using the following gradient: 5% B (0 min) −40% B (40 min) −55% B (45 min) −90% B (48 min) −90% B (54 min) −5% B (60 min). The fractions were collected every 1 min and consolidated into 10 fractions with equal intervals. The fractions were dried in a SpeedVac at 45°C and stored at 4°C until the analysis.

### 2.4 nLC−MS/MS Analyses

All samples above were resuspended with 0.1% formic acid (FA). NanoRPLC experiments were performed on an an EASY-nLC 1200 system (Thermo, CA, USA). Peptides were separated on a homemade C18 columns (5 μm, 100 Å, 150 mm×2.1 mm i.d., Durashell, Tianjin, China) with mobile phases (solvent A: 100% H_2_O and 0.1% FA; solvent B: 20% H_2_O and 80% ACN + 0.1% FA). The separation gradient was achieved by applying 8−30% B for 95 min, and 30−50% B for 25 min. An Orbitrap Exploris 480 (Thermo, CA, USA) mass spectrometer with a FAIMS Pro Interface (San Jose, CA, USA) was employed for MS/MS analysis. The electrospray voltage was set to 2.2 kV and the temperature of ion trans fer tube was set to 320 °C. Default settings were used for FAIMS with a total carrier gas flow of 4 L/min and compensation voltages (CV) of -45V and -65V. For both DDA and DIA analysis, the same LC and FAIMS settings were used for retention time stability. DDA were performed with full scan MS spectra acquired at a resolution of 60,000 fwhm (350−1500 m/z) with a normalized AGC target of 300% over a maximum of 20 ms. A 1.6 m/z isolation window and a fixed first mass of 110 m/z was used for MS/MS scans. Fragmentation of precursor ions was performed by higher energy C-trap dissociation (HCD) with 30% collision energies. MS/MS scans were performed at a resolution of 15,000 at m/z 200 with a normalized AGC target of 75% and a maximum injection time of 30 ms. Dynamic exclusion was set to 30 s to avoid repeated sequencing of identical peptides. DIA runs were carried out with the resolution set at 60,000 (at m/z 400) for full MS scans with the normalized AGC target of 300%, and the maximum injection time was set to 20 ms. The HCD collision energy was 30% and the MS/MS scans were performed at the resolution of 30,000 with the standard AGC target and auto maximum injection time. The mass list table was showed in Supplementary information 1.

### 2.5 MS Data Analysis

All samples acquired by DDA-MS were searched using Mascot node integrated within the Proteome Discoverer software (PD, version 2.5) configured with the human proteome database (Proteome ID: UP000005640; downloaded in 08/10/20 containing 38007 reviewed sequences). Common contaminants were added to the databases. Search parameters assumed tryptic digestion of peptides, a fragment ion mass tolerance of 20 ppm and a parent ion tolerance of 10 ppm. Carbamidomethylation (C) (+57.021 Da) and peptide N-terminal and lysine dimethyl labeling (+32.05641 Da (32)) were set as fixed modification, while oxidation (M) (+15.995 Da) set as variable modifications. False discovery rate (FDR) of identified peptides and protein were validated by the Percolator algorithm at 1% based on q-values.

The DIA-MS result were first converted to profile ms1 and ms2 using ProteoWizard^[16]^ v.3.0.6002. Then, by using self-made software, the spectral information such as m/z, retention time (RT), and ion intensity in the matching spectral library were output in mgf format. The mgf data were searched against the human fasta database (Proteome ID: UP000005640; Downloaded in 08/10/20 containing 38007 reviewed sequences) using Mascot node integrated within the Proteome Discoverer software (PD, version 2.5). Common contaminants were added to the databases. The precursor and fragment mass tolerances were set to 10 ppm and 20 ppm, respectively. A maximum of two missed cleavages was allowed for trypsin digestion. Carbamidomethylation (C) (+57.021 Da) and peptide N-terminal and lysine dimethyl labeling (+28.03130 Da (28), +30.04385 Da (30), +32.05641 Da (32), +34.06896 Da (34), +36.07567 Da (36)) were set as fixed modification, and the mixed five labeled channel were searched separately, together with oxidation (M) (+15.995 Da) set as variable modifications. False discovery rate (FDR) of peptide spectrum matches (PSMs) and identified peptides were validated by the Percolator algorithm at 1% based on q-values. Quantitation analysis was simultaneously performed by the self-made software, the intensities of b and y ion clusters of all PSMs were extracted within 2 mDa mass tolerance and used for relative quantitation. Peptide and protein ratios were calculated as the sum of the ion intensity matching the peptide and all quantified peptides matching the protein, respectively.

## 3 Results and discussion

### 3.1 Development of UV-assisted protein digestion (UVAPD), 5-plex labeling and MS-based high abundance protein removal quantification method

In clinical research, a large amount of analysis samples is often required, which requires a high-throughput, robust, and highly reproducible workflow. Here our goal is to develop a convenient workflow, from sample preparation to MS analysis, which can be used in a clinical context. Therefore, it should minimize all preparation and analysis steps while still accurately quantifying proteins of clinical interests. With this in mind, we decided to omit high-abundance plasma protein depletion, instead of performing a MS-based high abundance protein removal based 5-plex DIA analysis after rapid sample preparation.

For sample preparation, a new Hi-UV irradiation instrument was designed to accelerate protein digestion, which was composed of a high-energy density LED array and a sample condensation system. Based on such an instrument, an integrated workflow with combination protein denaturation, reduction and digestion was established (Figure 1). Herein, although by traditional in-solution sample preparation protocols, cysteine alkylation after protein reduction is an indispensable step, we ignored the cysteine alkylation step in our workflow since all sample preparation procedures were completed in an enclosed tube. Due to the good compatibility of the peptide buffer with MS detection, the resulting peptides from UVAPD were entered directly into an LC-MS/MS for further analysis without any additional peptide clean-up. The entire procedure took about 20 min and can readily be performed in a 96-well format and automated in a liquid handling platform, if desired.

**Figure 1.**
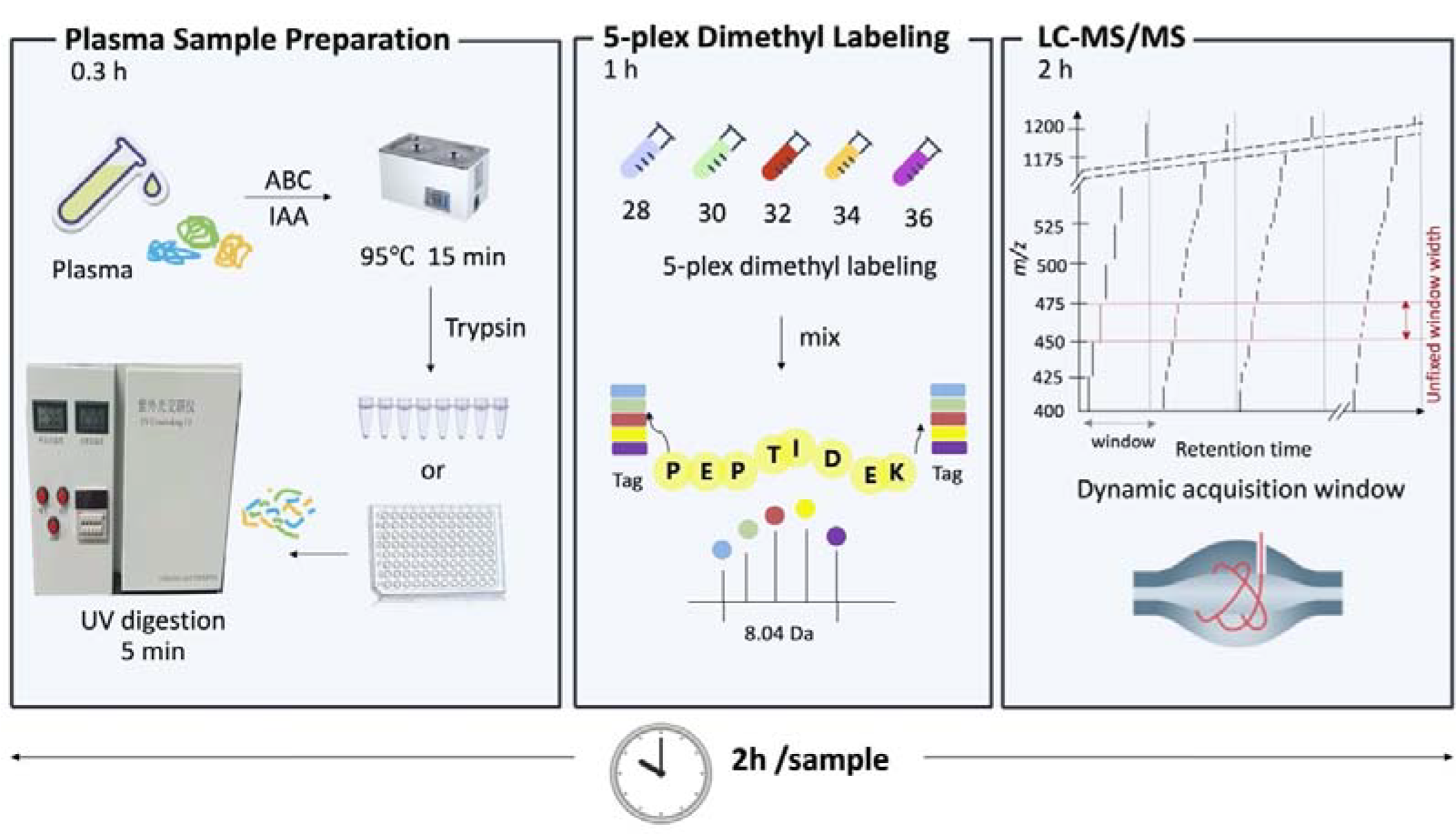
Scheme of UVAPD, 5-plex labeling and MS-based high abundance protein removal quantification method.

The prepared peptides were labeled with 5-plex dimethyl reagents and directly analyzed by LC-MS without the need for extra high abundance protein removal steps. Based on the advantages of fast, reliable, cost-efficient, and easily automated labeling^[17]^, together with recently been proved to be suited for multiplexed acquisitions of complex proteomes in DIA mode^[18]^, we labeled tryptic peptides with dimethyl reagents and obtained the peptide which N-terminal and lysine added with the mass tag of 28.0313 Da (28), 30.04385 Da (30), 32.0564 Da (32), 34.06896 Da (34), or 36.0757 Da (36). This increased the analysis throughput to 5-plex, which would realize analyze more plasma samples at the same time.

Then the DIA acquisition was divided into several time windows, each of which filtered out the mass range of high abundance peptides with retention time within this range. The time window should be set in an appropriate range, ensuring that the high abundance peptides that appeared during this period were completely removed, as they usually lasted for a long time. Besides, the numbers of removed high abundance peptides were needed to be taken into account. The more the peptides number removed, the smaller the mass range could be acquired. The less the peptides number removed, the incomplete removal effect. Therefore, it is necessary to find a balance between them. Thus, a convenient workflow came up to fasten the sample preparation process and improve proteome coverage which was promising for application in clinical research.

### 3.2 Evaluation on the performance of UVAPD

With human plasma as the sample, we firstly investigated the performance of UV irradiation to improve the efficiency of protein digestion. As shown in Fig. 2a, compared to traditional in-solution protocol, it could be seen that comparable number of protein identifications (393±32 vs. 385±12, n=3) could be achieved by tryptic digestion under 365 nm UV irradiation for 5 min, which was at least 100 times shorter than that (12 h) taken by traditional in-solution digestion.

**Figure 2.**
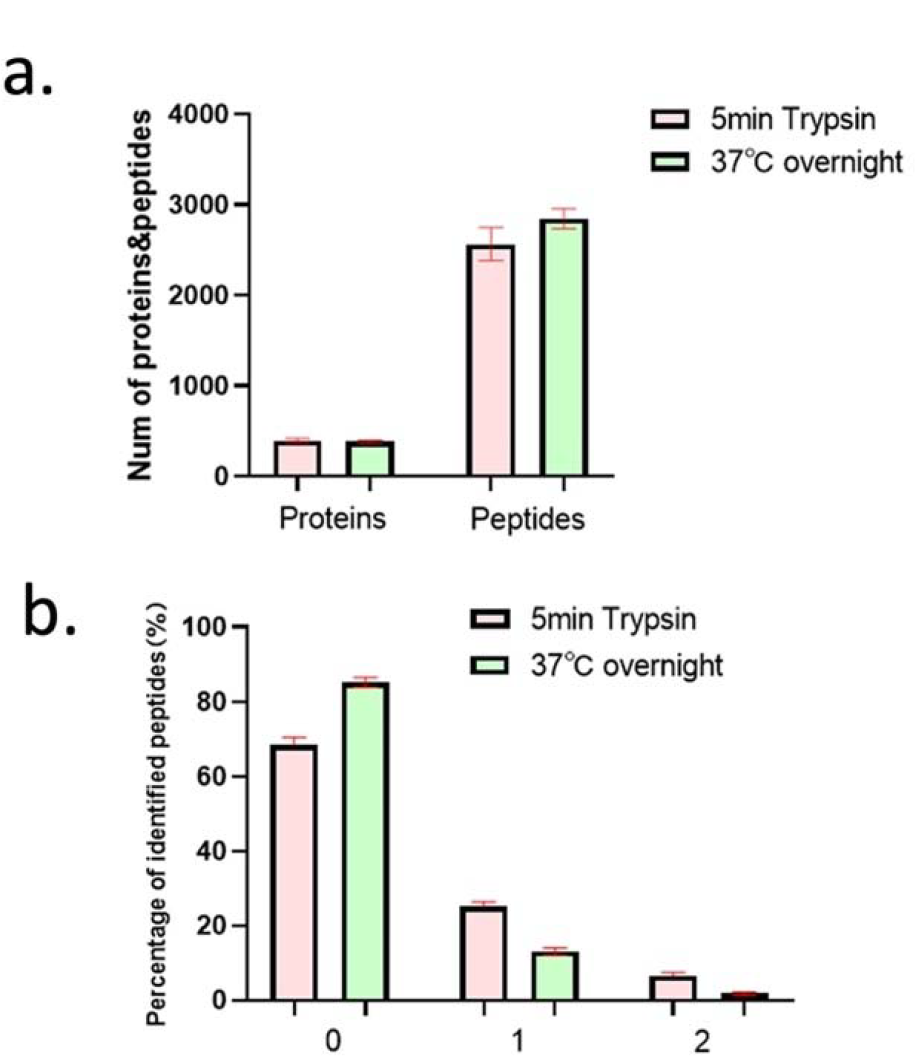
Comparison of UVAPD and traditional in-solution protocol. a, number of identified proteins and peptides. b, miss-cleavage rate.

However, it was also worthy noted that the miss-cleavage rate produced by UV-assisted digestion was much higher than that achieved by in-solution overnight digestion (Figure 2b), which might affect the accuracy of proteome quantification. Interestingly, we further combined trypsin with Lys-C in a mixed ratio of 1:1 (w/w) for protein digestion, although the number of HeLa protein identifications (4804 vs. 4787) were comparable, the missed cleavage rate (26% vs. 32%) was dramatically decreased (Table 1). We supposed that the complementarity of cleavage contributed by different specific enzymes significantly improved the efficiency of protein digestion. Using these optimal digestion conditions, the whole sample preparation time including protein denaturation, reduction, and digestion could be greatly shortened to about 20 min, which promises to significantly increase the throughput of large-cohort sample preparation.

**Table 1.** Comparison of different UV–assisted protein digestion strategies.

|  | Protein IDs | Peptide IDs | Miss-cleavage sites |  |  |
| --- | --- | --- | --- | --- | --- |
|  |  |  | 0 | 1 | 2 |
| 5min_trypsin_1 | 5351 | 31738 | 67.44% | 26.31% | 6.25% |
| 5min_trypsin_2 | 4588 | 25179 | 69.23% | 24.99% | 5.78% |
| 5min_trypsin_3 | 4473 | 24294 | 69.25% | 24.78% | 5.97% |
| 5min_Trypsin+lys-C_1 | 5230 | 31210 | 69.11% | 25.2% | 5.69% |
| 5min_Trypsin+lys-C_2 | 4605 | 24861 | 77.33% | 18.9% | 3.76% |
| 5min_Trypsin+lys-C_3 | 4528 | 24037 | 77.19% | 19.03% | 3.78% |

### 3.3 Quantitation depth and accuracy of 5-plex labeling and MS-based high abundance protein removal quantification method

Plasma contains a large amount of HAPs, and the “masking” effect caused by these HAPs makes it challenging to detect LAPs such as cytokines and other essential proteins that are typically present at levels below sub ng/mL^[19]^. Considering the non-specific nature of existing methods for removing HAPs, as well as the high cost and limited sample loading capacity^[20]^, we chose to filter the MS acquisition window of HAPs to achieve specific and efficient removal. Meanwhile, by combining DIA-based 5-plex labeling method, achieved unbiased and high-throughput plasma proteome analysis.

We first optimized how to filter the acquisition window for HAPs. Firstly, it was necessary to obtain the m/z and retention time (RT) information of the peptides that needed to be removed. Then, these peptides were sorted by RT, and high abundance peptides that appeared during a certain period of time were removed. For example, if there were 10 peptides needed to be removed within the first five minutes, the mass range of these peptides would no longer be acquired within 0-5 min, and the subsequent acquisition would also be similar. We first conducted DDA analysis on plasma to select peptide of 2-3 charge form from the TOP14 protein for removal. Table 2 showed the proportion of high abundance peptides with different numbers to the total intensity, with 50 peptides accounting for 34.69%, 100 peptides accounting for 47.83%, 200 peptides accounting for 59.80%, and 500 peptides accounting for 67.50%. And the peptide details were showed in Supplementary information 2. We chose these numbers of peptides for removal as for their intensity varied in a stepped pattern. Next, we divided these peptides into 5-minute or 10-minute window based on retention time to obtain a suitable window for more complete removal of high abundance proteins. Therefore, we obtained 8 combinations, including sequentially removing 50, 100, 200, or 500 peptides within 5 or 10 minutes, to obtain the best window for deep plasma proteome analysis.

**Table 2.** Ratio of intensity to total intensity of different numbers of high abundance peptides.

| Peptide Number | 50 | 100 | 200 | 300 | 500 | all |
| --- | --- | --- | --- | --- | --- | --- |
| Intensity | 3.58e<sup>11</sup> | 4.94e<sup>11</sup> | 6.17e<sup>11</sup> | 6.66e<sup>11</sup> | 6.96e<sup>11</sup> | 7.02e<sup>11</sup> |
| Percentage | 34.69% | 47.83% | 59.80% | 64.56% | 67.50% | 68.09% |

As shown in Table 3, the combination of removing 100 peptides with 10-minute window showed the deepest depth, quantifying 636 plasma proteins. However, the combination of removing 100 peptides with 5-minute window showed similar depth. Therefore, we compared the number of quantified proteins between the five channels under two conditions. The Venn diagram showed good reproducibility between the 1:1:1:1:1 mixed five labeled channels in the combination of removing 100 peptides with 10-minute window (Figure 3a). Also, the proteins quantified in the 10-minute window were queried for protein concentration through the PAXDB database, covering the maximum 9 orders of magnitude contained in the database (Figure 3b), and compared to the 5-minute window, most of the proteins quantified are of medium to low abundance. Besides, a large number of low abundance protein biomarkers approved by Food and Drug Administration (FDA)^[21]^ were present in this data set such as collagen alpha-1(P02452), myeloperoxidase (P05164), and epidermal growth factor receptor (P00533), showing good prospects for discovering plasma biomarkers in clinical research. We further evaluated the quantification accuracy, and the average relative error of quantification within the five channels was 6.9% (Figure 3c), indicating excellent quantitative performance. Therefore, the MS-based high abundance protein removed 5-plex DIA quantification method exhibited an in-depth and accurate quantification for plasma proteome, heralding great potential for providing a solution for the analysis of trace plasma proteome samples in clinical queues.

**Table 3.** Number of identification proteins with different removed peptide numbers in different time windows under five labeling channels.

|  | 5-minute window |  |  |  | 10-minute window |  |  |  |
| --- | --- | --- | --- | --- | --- | --- | --- | --- |
|  | Remove 50 peptide<br>s | Remove 100 peptide<br>s | Remove 200 peptide<br>s | Remove 500 peptide<br>s | Remove 50 peptide<br>s | Remove 100 peptide<br>s | Remove 200 peptide<br>s | Remove 500 peptide<br>s |
| 28 | 576 | 625 | 558 | 462 | 617 | 634 | 581 | 415 |
| 30 | 585 | 623 | 556 | 459 | 588 | 635 | 578 | 415 |
| 32 | 581 | 623 | 551 | 448 | 616 | 634 | 579 | 420 |
| 34 | 573 | 615 | 555 | 447 | 609 | 635 | 573 | 399 |
| 36 | 576 | 611 | 557 | 452 | 576 | 633 | 570 | 415 |
| Total | 615 | 634 | 599 | 531 | 628 | 636 | 607 | 501 |

**Figure 3.**
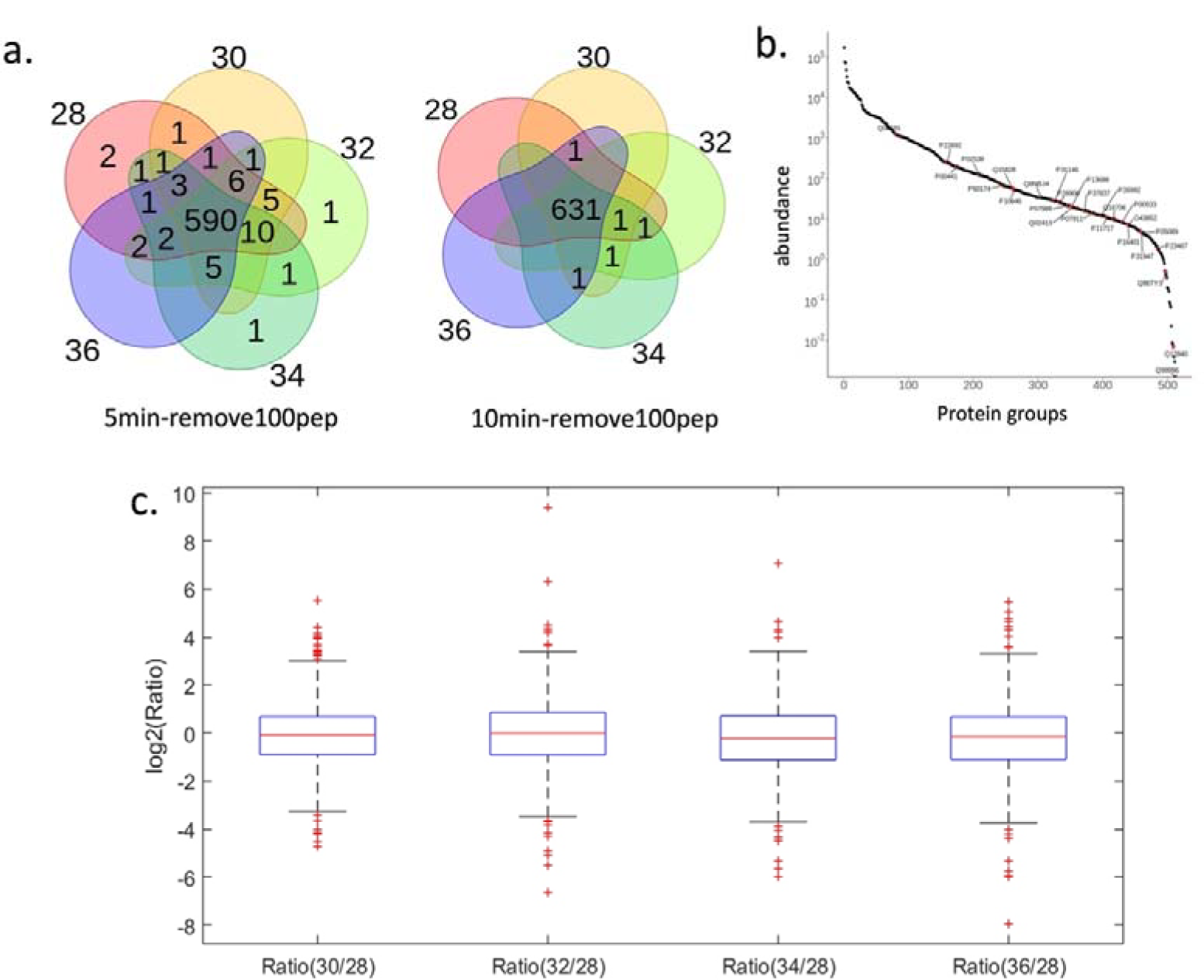
Depth and accuracy evaluation of MS-based high abundance protein removed 5-plex DIA quantification method. a, The venn diagram of identified proteins under five channels in two combinations. b, Distribution of plasma protein intensity. The red section shows the proteins only identified in the window of removing 100 peptides within 10 minutes compared to the window of removing 100 peptides within 5 minutes. c, Quantitation results of plasma proteins mixed at the ratio of 1:1:1:1:1. Box plots showed the median (centred red line), first and third quartiles (lower and upper box limits, respectively), 1.5 times the interquartile range (whiskers) and outliers (cross).

## 4. Conclusions

In summary, we developed UV-assisted protein digestion and MS-based high abundance protein removal based 5-plex DIA method to solve the difficulty of high abundance protein removal and limitation of high-throughput quantitative coverage in trace plasma samples. We achieved efficient enzymatic hydrolysis by UVAPD procedure in 20 minutes, and the depth was reached to traditional in-solution protocol. Also, high abundance proteins were deleted by MS-based high abundance protein removal method, and the acquisition method for removing 100 peptides with 10-minute window were determined, achieving the quantification of 636 plasma proteins with a dynamic range spanning 9 orders of magnitude. With 5-plex labeling, the mass spectrometry analysis time of each sample was reduced to 30 minutes, and achieved an accurate quantification with an average relative error of 6.9%. This process provides a technical protocol for fast, simple, and deep coverage analysis of trace plasma samples.

## Acknowledgment

The authors gratefully acknowledge the financial supports from National Key Research and Development Program of China (2020YFE0202200, 2022YFC3341003), Innovation Program of Science and Research from Dalian Institute of Chemical Physics, Chinese Academy of Sciences (DICP I202143), and Applied Basic Research Program of Liaoning Province (2023JH2/101300126).

